# Visualizing the Spatio-Temporal Dynamics of Clonal Evolution with *LinG3D* software

**DOI:** 10.1101/2024.03.05.583631

**Authors:** Anjun Hu, Awino Maureiq E. Ojwang’, Kayode D. Olumoyin, Katarzyna A. Rejniak

## Abstract

Cancer clonal evolution, especially following anti-cancer treatments, depends on the locations of the mutated cells within the tumor tissue. Cells near the vessels, exposed to higher concentrations of drugs, will undergo a different evolutionary path than cells residing far from the vasculature in the areas of lower drug levels. However, classical representations of cell lineage trees do not account for this spatial component of emerging cancer clones. Here, we propose the *LinG3D* (Lineage Graphs in 3D) algorithms to trace clonal evolution in space and time. These are an open-source collection of routines (in MATLAB, Python, and R) that enables spatio-temporal visualization of clonal evolution in a two-dimensional tumor slice from computer simulations of the tumor evolution models. These routines draw traces of tumor clones in both time and space, with an option to include a projection of a selected microenvironmental factor, such as the drug or oxygen distribution within the tumor. The utility of *LinG3D* has been demonstrated through examples of simulated tumors with different number of clones and, additionally, in experimental colony growth assay. This routine package extends the classical lineage trees, that show cellular clone relationships in time, by adding the space component to show the locations of cellular clones within the 2D tumor tissue patch from computer simulations of tumor evolution models.

## Background

During tumor development, cells undergo somatic mutations that may offer them a survival advantage over other cells. This, in turn, can lead to the emergence of distinct cell populations, i.e., clones (1). Tumor clonal diversity may be especially important when anti-cancer treatments are applied, since distinct clones can be affected differently, leaving those with a survival advantage, such as resistance to the drugs or radiotherapy, to persist. Thus, understanding how individual tumor clones emerge and spread may be vital in designing effective treatment therapies.

There are several ways to graphically present tumor clonal evolution. In one routinely used method, called “jellyfish” graphs or tumor evolution graphs, the horizontal axis represents the time along which plots with different colors indicate distinct genotypes. The height of each plot corresponds to that genotype’s relative abundance. If the clones generate descendant genotypes, they are shown as new areas emerging from inside their parents. These graphs showing changes in the sizes or frequencies of the clones over time can be drawn using the *ggmuller* (2), *fishplot* (3), or *evofreq* (4) routines. Another way to illustrate the development of tumor evolution is to show cell lineages in the form of a phylogenetic tree. In this graph, one starts with an initial cell (tree root) and connects each mother cell with its two daughter cells until all final descendants are reached (tree leaves). This produces a binary tree in which time or cell generations are represented by the axis joining the tree root with tree leaves, and the tree nodes are located at the times of the mother cells’ division. These graphs showing the dynamics of clone development can be drawn using the *phytree* (5), *ggtree* (6), or *TreeDyn* (7) routines.

However, the tumor clonal composition, such as the sizes and diversities of clones, depends strongly on the microenvironment surrounding the tumor. For example, cells located near the vessels are exposed to higher levels of bloodborne therapeutics and thus can be more prone to drug-induced death, thus generating smaller cellular clones, unless the cells develop resistance to that treatment. Therefore, they can undergo a different evolutionary path than cells residing far from vasculature, in areas with low drug levels where cells can proliferate more frequently and die less often, thus producing larger cellular clones or clones with more mutations. This spatial resolution is lacking in the classical clonal graphs, as those described above. We have designed the *LinG3D* routines by incorporating the space component to show the locations of cellular clones within the 2D patch of the tumor tissue, that is in relation to other cells, vasculature, or the distribution of drugs or nutrients. These routines are intended to visualize the 3D lineage trees for data generated by computational models of tumor evolution. However, they can be applied to experimental data, if appropriate information can be collected in the laboratory. In this paper, we describe four routines that form the *LinG3D* package allowing to visualize (i) an individual clone with all generated cells, (ii) an individual clone with no dead branches, i.e., the clone including only alive cells, (iii) all clones with all generated cells, and (iv) all clones with no dead branches. The utility of these routines is demonstrated using two examples of simulated tumors with different numbers of clones and an example of an experimental colony growth assay.

### Design and Implementation

*LinG3D* has been created as a suite of routines that enable visualization of the spatio-temporal evolution of the cellular clones arising during computational simulations of mathematical models of tumor growth and/or its response to treatments. These routines display not only the mother cell-daughter cell linear relationship as in a classical lineage tree, but also show which subspace of the whole tumor tissue each clone is occupying during its development and how these locations change over time. Four independent routines are provided to draw either an individual clone or all clones together, and with either all cells generated during the time of the computer simulation or the cells whose successors survived to the end of the simulation (i.e., by omitting branches with cells that died or left the simulation domain). The architecture of the four routines included in the *LinG3D* package is shown in Fig1.

**Fig. 1.**
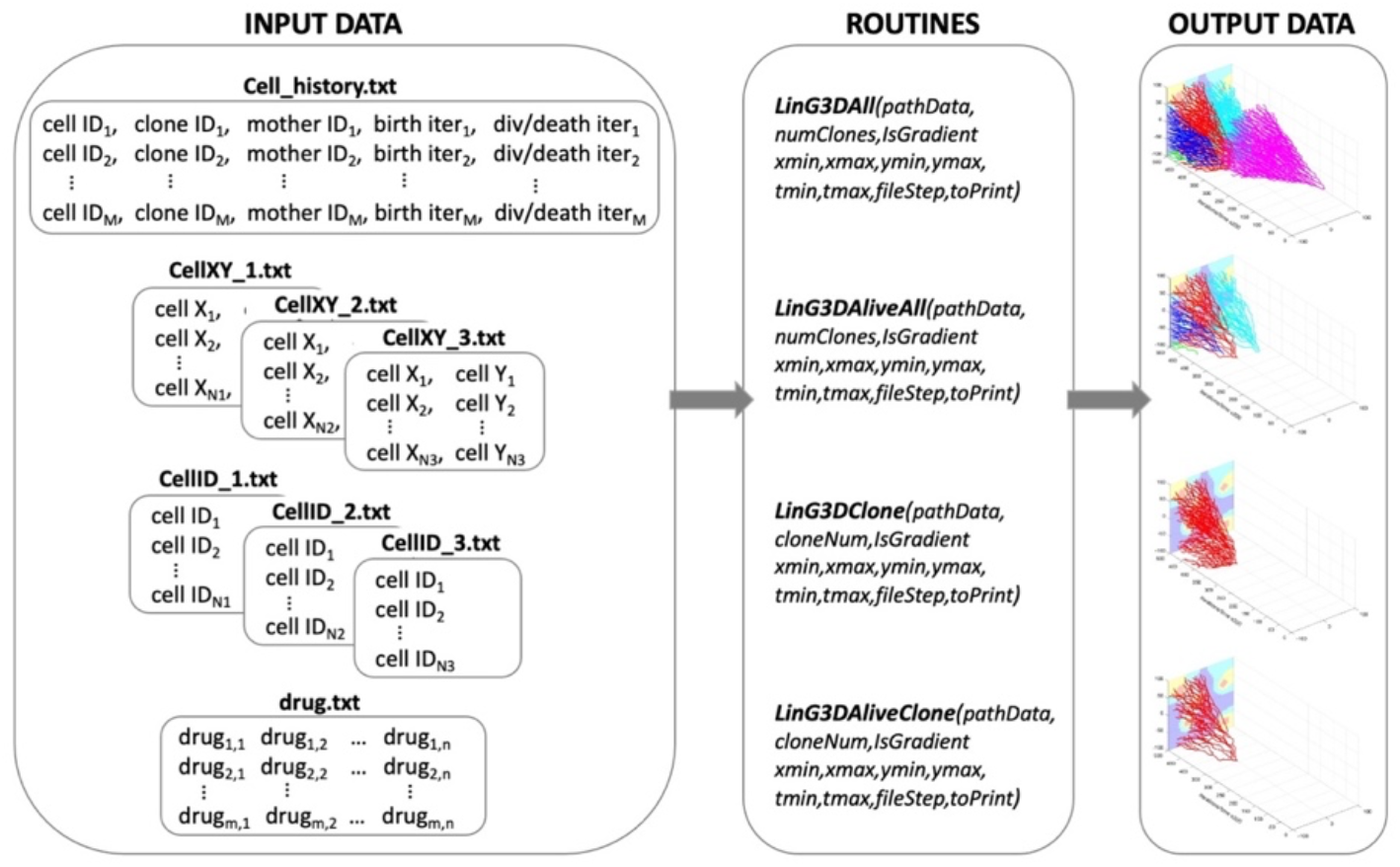
Architecture of the *LinG3D* routine package. The specified four types of text files are required as the input data all routines within the LinG3D package; each routine generates a figure as an output data.

The following routines are included in the *LinG3D* package.

### LinG3DAll

This routine draws a 3D spatio-temporal lineage tree taking into account all cells from all cellular clones. To do so, the routine starts with an initial cell (the tree root), and draws branches of the lineage tree for all mother-daughter cell pairs (straight line segments connecting these two cell positions), until the cells of each clone are included and the terminal cells with no descendants (clone leaves) are reached. A full 3D lineage tree with five clones is shown in Fig.2A.

**Fig. 2.**
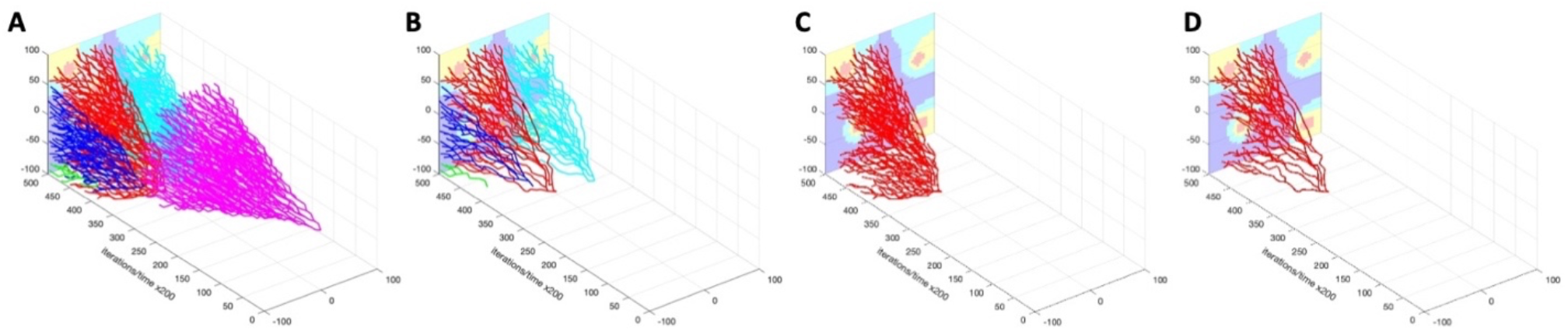
Examples of outcomes from four *LinG3D* routines. **A:** a 3D full lineage tree with all cells and clones (*LinG3DAll* routine); **B:** a 3D lineage tree with all surviving cells (*LinG3DAliveAll* routine); **C:** a 3D lineage tree with all cells for a single clone (*LinG3DClone* routine); **D:** a 3D lineage tree with all surviving cells in one clone (*LinG3DAliveClone* routine). In all examples, the drug distribution is shown in the background with concentrations from high to low represented by: red-yellow-cyan-blue.

### LinG3DAliveAll

This routine visualizes a 3D spatio-temporal lineage tree for all cellular clones, but includes only those cells for which at least one successor cell survived until the end of the simulation. Therefore, it omits branches with cells that died or that left the computational domain. The routine starts with cells in the last simulated iteration (tree leaves) and traces back all branches of the lineage tree for all daughter-mother cell pairs until the initial cell of the clone (the root) is reached. Fig.2B shows the 3D lineage tree of alive cells from the full lineage tree from Fig.2A.

### LinG3DClone

This routine visualizes a 3D spatio-temporal lineage tree for one selected cellular clone showing all cells in that clone. This routine starts with the initial cell of the selected clone (the clone tree root) and draws all mother cell-daughter cell branches of the lineage tree for cells belonging to that clone until all cells are included and the terminal cells with no descendants (clone leaves) are reached. Fig.2C shows the full 3D lineage tree for one clone from Fig.2A.

### LinG3DAliveClone

This routine draws a 3D spatio-temporal lineage tree for the surviving cells in the selected clone. The routine starts with the last iteration and selects cells that belong to the given clone (clone leaves). Next, it draws the branches for all daughter-mother cell pairs until the initial cell of that clone (the clone root) is reached. Fig.2D shows the 3D lineage tree for alive cells from one clone from Fig.2B.

### Required format of the input data

All input data required by ***LinG3D*** routines consists of text files of four kinds. One text file with information about the history of all cells to trace a given cell’s predecessors or successors, i.e., cell’s index, the index of a clone it belongs to, the index of its mother cell, as well as the iteration numbers at which the cell was born, and at which it either divided or died. Moreover, for every iteration saved, two text files are required, one containing indices of all cells alive at this iteration, and the other with coordinates of cells present at this iteration. If the background gradient of drug or metabolite should be drawn (*IsGradient*=1), then a text file with a matrix of drug concentrations must be provided. Each of these text files is described in detail below.

The cell history data contains one cell per row (file in our code: *cell_history.txt*):

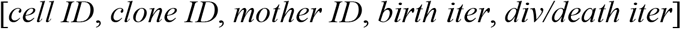

where columns contain *cell ID*—a unique ID number for the cell, *clone ID*—a number unique to a given clone to which the cell belongs, *mother ID*—a unique ID number of the cell!s mother cell, *birth iter*—the iteration number at which the cell was born, and *div/death iter*— the iteration number at which the cell either divided into two daughter cells or died. If the cell has not divided and is still alive, this element is equal to 0.

Moreover, for each iteration at which the lineage tree branches are drawn (denoted by *_#*), the following files are required to contain the x and y coordinates of all cells present at that iteration, one cell per row (file in our code: *cellXY_#.txt*):

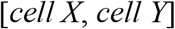

and the corresponding unique cell ID numbers, one cell per row (file in our code: *cellID_#.txt*):

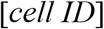

For every iteration saved (*_#*), the order of cells in these two files must be identical.

Additionally, if the microenvironmental factor distribution (such as, oxygen or a drug) is to be drawn in the background, the file containing the factor concentrations per grid point should be saved as a 2D matrix (file in our code: *drug.txt*).

### Expected output data

The output data generated by each of the ***LinG3D*** routines is an on-screen figure with a corresponding 3D lineage tree graph (Fig.2A-D) showing clonal expansion in space (cell locations within the tumor tissue in the xz-plane) and in time (tumor clone’s temporal evolution along the y-axis). This figure can be automatically saved as a graphical file in the jpeg format, if *toPrint*=1.

### User-specified input function arguments

Each ***LinG3D*** routine requires that values of several input function arguments are specified by the user to represent their own simulation setup:

1. *pathData* – a text argument that defines the name of the main directory in which the subdirectory /data/ must be located with all input data files.
2. *numClones or cloneNum* – a natural number argument that specifies either the total number of clones to be drawn (*numClones* in routines: ***linG3DAll*** and ***linG3DAliveAll***), or a specific clone number that will be drawn (*cloneNum* in routines: ***linG3DClone*** and ***linG3DAliveClone***); the clones are numbered starting at 0.
3. *xmin, xmax, ymin*, and *ymax* – four real number arguments that specify the domain boundaries [x_*min*_, x_*max*_] × [y_*min*_, y_*max*_] used to define a rectangular patch of a tissue. *tmin* and *tmax* –
4. two real number arguments that specify the initial and final iteration numbers. Note, that these numbers are used to optimize the spacing along the y-axis that represents the number of recorded data (*timeStep* variable). This is done for visualization purposes, such that all three axes have comparable scale.
5. *fileStep* – a natural number argument that specifies the progression step indicating frequency of the data to be sampled for the 3D lineage three visualization. Note that, more frequent data sampling will show more details of the simulation: smaller changes in cell locations between time points at which the branches are drawn, that results in more tortuous tree branches (Fig.3A). On the other hand, less frequent data sampling for drawing results in more linear tree branches as some intermediate cell positions will be omitted (Fig.3B).
6. *IsGradient* – a binary number argument that indicates whether to draw the background concentration of a specific environmental factor or not: 1 to draw (Fig.3B), 0 not to draw (Fig.3C). The spatial distribution of the microenvironmental factor, such as oxygen or drug, is projected on the xz-plane. Note, that the provided concentration values (min and max drug levels) are used to optimize the split of concentration values into four groups, in order to draw each group in a different color (from the highest concentration to lowest: red— yellow—cyan—blue). This is done for visualization purposes and allows to use four distinct colors to draw the background gradient.
7. *to Print* – a binary number argument that indicates whether to save the final figure or not (1 to save, 0 not to save).

**Fig. 3.**
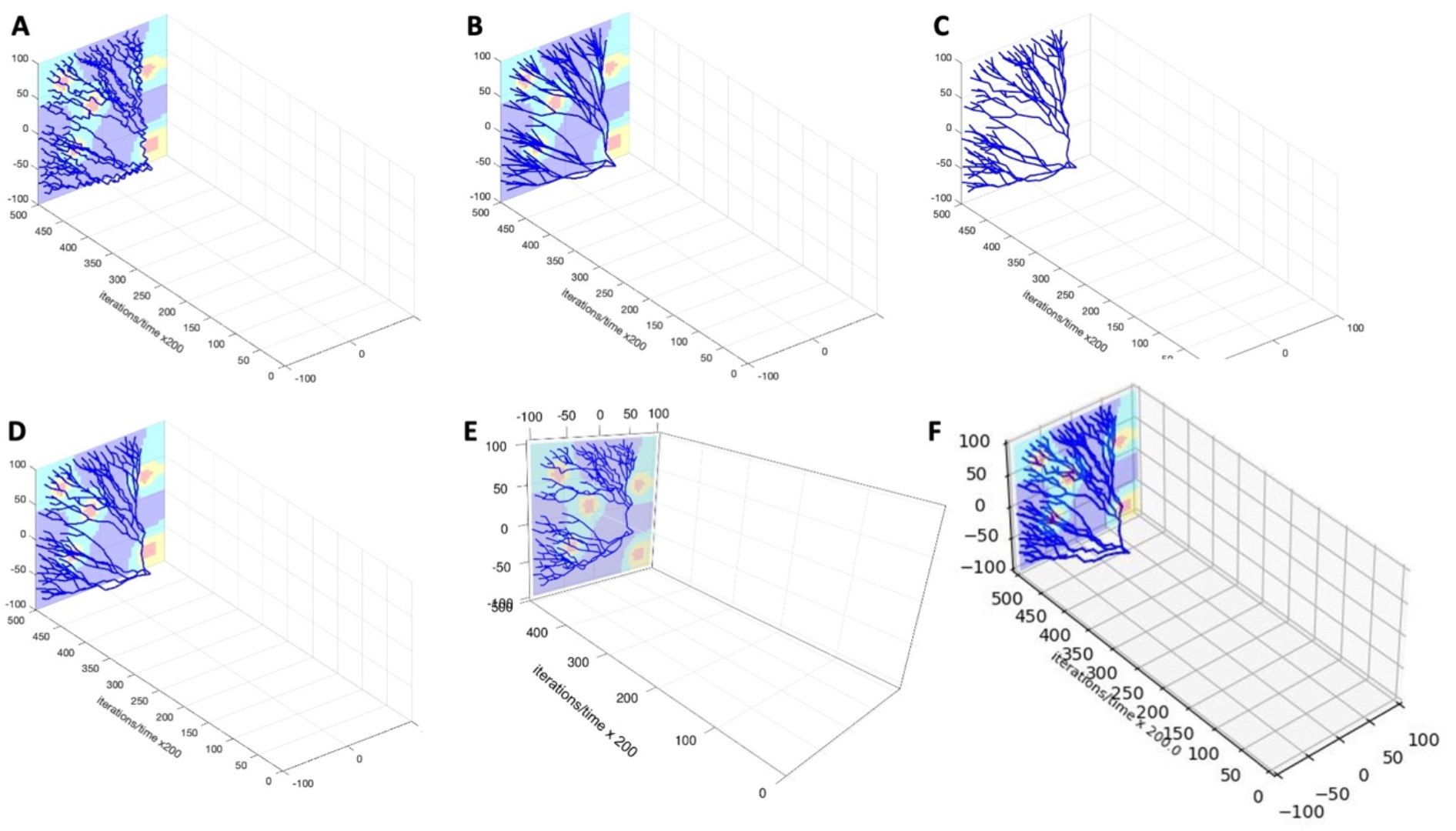
User-specified parameters and computational language options. **A:** A 3D lineage tree of surviving cells for one clone (*LinG3DAliveClone*) drawn with data sampling parameter *fileStep*=250; **B:** The same lineage tree drawn with data sampling parameter *fileStep*=5000; **C:** The same lineage tree drawn with no background *IsGradient*=0 and with data sampling parameter *fileStep*=2500. **D-F:** 3D lineage trees of the same single clone of surviving cells drawn with routines implemented in MATLAB (**D**), R (**E**), and Python (**F**), respectively (*fileStep*=2000).

This implementation is unified across the MATLAB, Python, and R routines.

### Available routine versions

The *LinG3D* routines have been implemented in different computational systems and each of the discussed routines is provided in three computational languages: MATLAB (Fig.3D), R (Fig.3E), and Python (Fig.3F). The following additional libraries are required to run the routines in R: *rgl* for 3D visualization (8), *rapportools* for vector and matrix operations (9), and *readr* for reading external files (10), and *devtools* for GitHub installation (11); MATLAB and Python routines do not require installation of any non-standard toolboxes or libraries.

## Results

To illustrate the results produced by each routine in the *LinG3D* package, we applied them to two simulated tumors that differ in the number of generated clones and one experimental colony growth assay with previously identified cellular clones. In both simulated cases, a growing colony of tumor cells is exposed to a cytotoxic drug, resulting in massive cell death. However, the acquired mutations make cells drug-resistant and allow for the emerging cellular clones to persist. These two examples vary only in one parameter: the probability of cell mutation which results in very different numbers of the generated cellular clones. In the experimental colony growth assay, the cells are grown in a drug-free medium in a standard well, where they can freely move and proliferate. Over the course of the experiment, the cells were able to divide a small number of times leading to the emergence of several small clones.

### Case study of a simulated tumor with a small number of clones

In this computational model, a small patch of the tissue with five nonuniformly spaced vessels is used. A single tumor cell starts proliferating until a cell colony of a noticeable size is established. Then, the intravenously injected drug diffuses out of the vessels forming an irregular gradient within the tissue. All cells absorb the drug from their vicinity and can die if the accumulated drug level is high. Upon cell division, the daughter cells can mutate and the mutated cells acquire drug resistance giving rise to a mutated drug-resistant cellular clone. In this example, the probability of cell mutation is low (𝒫^*mut*^=0.005), thus, a small number (9) of cellular clones is generated. In Fig.4, six snapshots from this simulation are shown. Starting from the original mother cell (Fig.4A), a small clone of non-mutated cells has developed (Fig.4B,C). Since the probability of mutation is small, only two mutated cells emerged before the drug was injected (Fig.4D). Extensive cell death is observed upon drug administration and its diffusion throughout the tissue

**Fig. 4.**
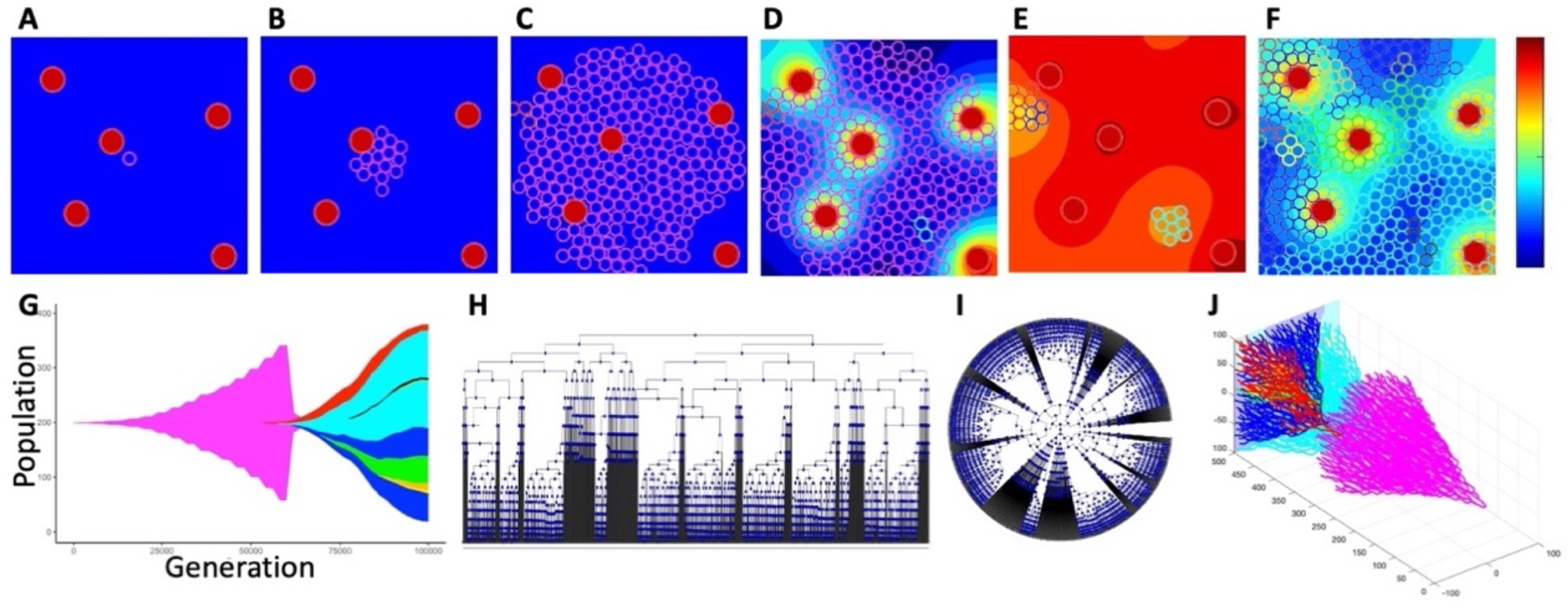
Evolution of a 9-clone tumor. Tumor growth simulated using *tumorGrowth_example1* with a parameter 𝒫^*mut*^=0.005: **A**. an initial tumor cell (pink, iteration 0); **B**. a growing cluster of non-mutated tumor cells (pink, iteration 18,000); **C**. the emergence of the first mutated cell (red, top left edge, iteration 56,500); **D**. the beginning of drug injection (second mutated clone in cyan, iteration 60,250); **E**. drug-induced cell death (iteration 63,250); **F**. the final configuration with 9 cellular clones (iteration 100,000). The color bar represents the drug concentration. **G**. the corresponding tumor jellyfish evolution graph (*ggmuller* routine (2)); Clone colors correspond to cell colors in **A-F. H-I**. a classical lineage tree showing the relationship between mother and daughter cells in square and radial configuration (*phytree* routine (5)); **J**. the corresponding full 3D lineage tree (our *LinG3DAll* routine).

(Fig.4E). The surviving cells have now more space to proliferate as the overcrowding is no longer an issue. However, the number of clones remains small (9 at the end of the simulation) since the mutation probability is low. All mutated cells are drug-resistant which leads to a repopulation of the whole tissue (Fig.4F). The details of this mathematical model are provided in the Appendix. The corresponding tumor jellyfish evolution graph (Fig.4G) shows changes in clonal abundance (population) over time (generation). The color of each clone corresponds to colors of cells in Fig.4A-F. The classical phylogenetic tree, in which tree branches connect nodes representing a mother and their two daughter cells, and with the branch length corresponding to the evolution time, is shown in Fig.4H in the horizontal configuration and in Fig.4I in the radial configuration. The full 3D lineage tree is shown in Fig.4J.

To obtain a better understanding of the tumor spatio-temporal evolution, we inspected the 3D lineage trees of all clones together and of the three most prominent individual clones (out of 9) from this simulation, including the initial clone. Fig.5 (drawn with R-based routines) shows pairs of 3D lineage trees: trees that include traces of all cells belonging to a given clone (Fig.5A-D) and trees that include only traces of cells that survived to the end of the simulation (Fig.5A!-D!). Each clone is denoted by a different color. In all cases, the trees with alive cells are smaller than the trees with all cells, since many cells have died (due to random cell death or drug-induced cell death) or have left the computational domain. In some cases, there are no surviving cells—the most obvious example is the initial clone of drug-sensitive cells (clone #0, Fig.5B!) for which all cells have died after the drug was administered.

**Fig. 5.**
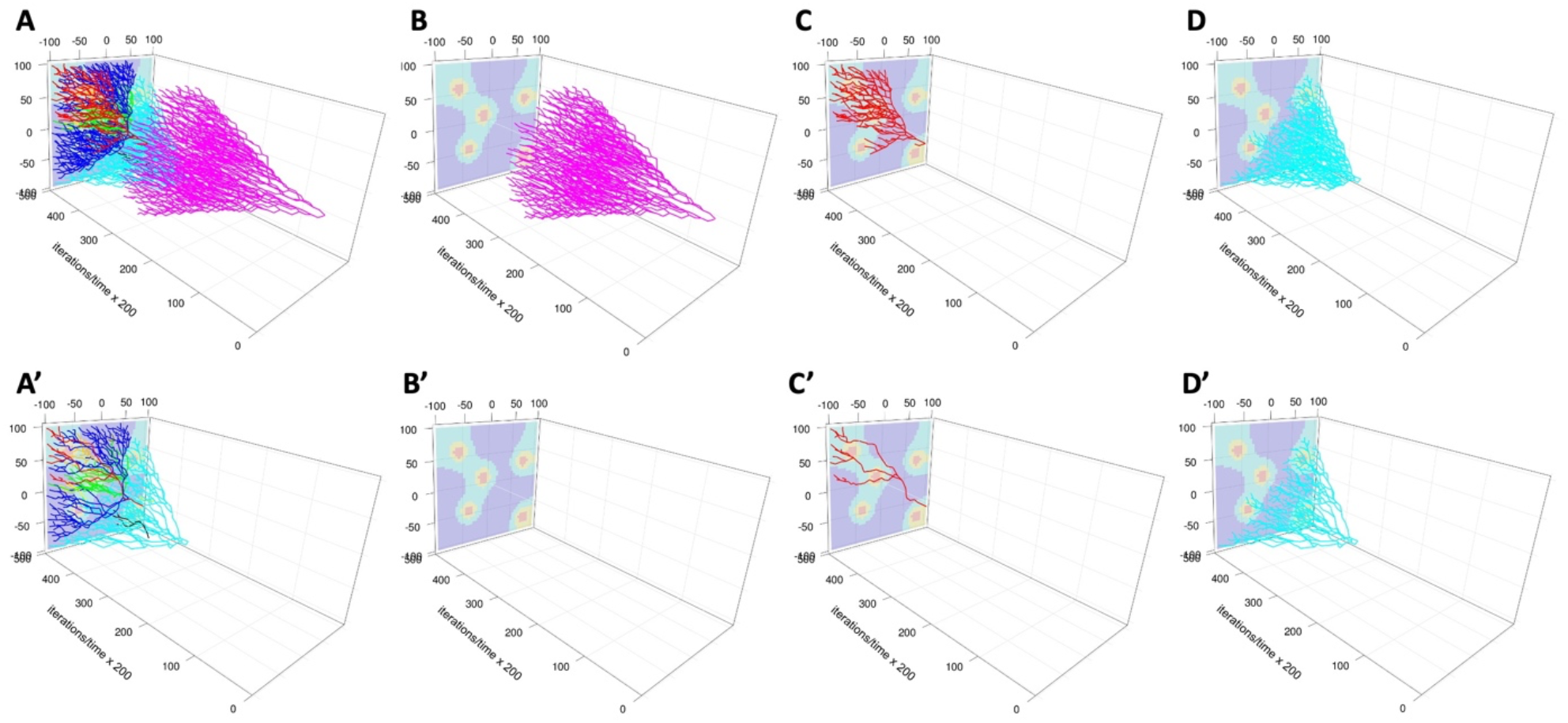
3D lineage trees of individual clones drawn with R routines. For each clone (denoted by a different color), the top row shows the 3D lineage tree with all cells belonging to that clone (**A** *LinG3DAll* and **B-D** *LinG3DClone* routines), while the bottom row includes only the cells that survived to the end of the simulation (**A’** *LinG3DAliveAll* and **B’-D’** *LinG3DAliveClone* routines). **B-B’:** initial clone #0; **C-C’:** mutated clone #1; **D-D’:** mutated clone #2.

To generate these 3D lineage trees, we had to specify the following function arguments: data directory (`pathData=‘exampleB005’`), spatial domain (`xmin=-100 xmax=100 ymin=-100 ymax=100`), temporal domain (`tmin=0 tmax=10000`), frequency of sampled data (`fileStep=250`), the background indicator (`IsGradient=1`), and the figure saving indicator (`toPrint=1`). For drawing all clones, the total number of clones (`numClones=9`) was specified. For drawing individual clones, the clone numbers (`cloneNum=0,1,2`) were used, respectively. The presented 3D lineage trees show that the initial clone develops and spreads uniformly in space before the drug is injected (Fig.5B) and then, upon drug diffusion, all these drug-sensitive cells die (Fig.5B!). However, the two presented drug-resistant clones are more localized and grow to occupy specific niches: the top-left tissue corner (Fig.5C) and the bottom-right corner (Fig.5D). This information is not available from classical jellyfish evolution graphs (Fig.4G) or phylogenic trees (Fig.4H,J).

### Case study of a simulated tumor with a large number of clones

In the second example, the probability of cell mutation is higher (𝒫^*mut*^=0.05) resulting in a large number (147) of cellular clones that are also more dispersed within the tumor tissue when compared to the previous example. In Fig.6, six snapshots from this simulation are shown. Like in the previous case, we start with one original mother cell (Fig.6A) that generates a small clone of non-mutated cells (Fig.6B). The growing tumor cell cluster is similar in shape to that in the previous example because we fixed the seed of the random number generator. However, since the probability of mutation is larger here, fifteen cells have mutated before the drug was administered (Fig.6C,D). Drug injection and its intratumor diffusion lead again to a massive cell death (Fig.6E) that allows more cells to proliferate as the cells are no longer overcrowded. More cell divisions also lead to an increase in the number of cell mutations (147 at the end of the simulation). All mutated cells are drug-resistant which leads to a repopulation of the whole tissue (Fig.6F). The corresponding tumor evolution graph (Fig.6G) shows changes in clonal abundance (population) over time (generation). The color of each clone corresponds to colors of cells in Fig.6A-E. The classical phylogenetic tree in the horizontal configuration is shown in Fig.6H and in the radial configuration in Fig.6I. The full 3D lineage tree is shown in Fig.6J.

**Fig. 6.**
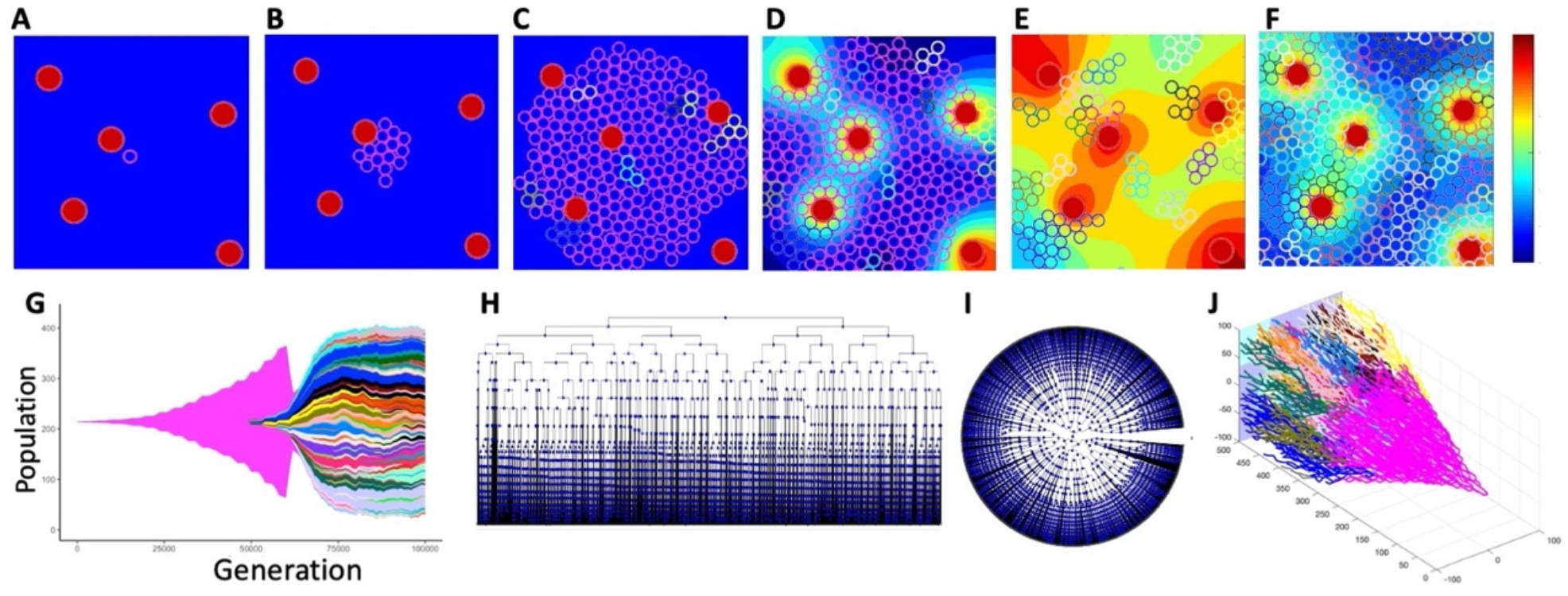
Evolution of a 147-clone tumor. Tumor growth simulated using *tumorGrowth_example2* with a parameter 𝒫^*mut*^=0.05: **A**. an initial tumor cell (pink, iteration 0); **B**. a growing cluster of non-mutated tumor cells (pink, iteration 18,000); **C**. ten mutated cells have already emerged (multiple colors, iteration 56,500); **D**. the beginning of drug injection (iteration 60,250); **E**. drug-induced cell death (iteration 63,250); **F**. the final configuration with 147 cellular clones (iteration 100,000); the color bar represents the drug concentration. **G**. the corresponding tumor evolution graph (*ggmuller* routine (2)); clone colors correspond to cell colors in **A-F. H**,**I**. classical lineage trees showing the relationship between mother and daughter cells in square and radial configurations (*phytree* routine (5)). **J**. the corresponding full 3D lineage tree (our *LinG3DAll* routine).

Since this simulation generated a large number of clones (147), we present here the 3D lineage trees for the three most prominent clones only. Fig.7 (drawn with Python-based routines) shows pairs of 3D lineage trees with the top row including traces of all cells belonging to a given clone (Fig.7A-D) and the bottom row including traces of cells that survived to the end of the simulation (Fig.7A’-D’). As in the previous case, each clone is denoted by a different color. These clones are smaller and much less dispersed than in Example 1 since the cells have mutated more often and generated more clones. Again, the trees with alive cells are smaller than the trees with all cells, because many cells have died (due to random cell death or drug-induced cell death) or have left the computational domain. In some cases, there are no surviving cells, including the initial drug-sensitive clone (Fig.7B’).

**Fig. 7.**
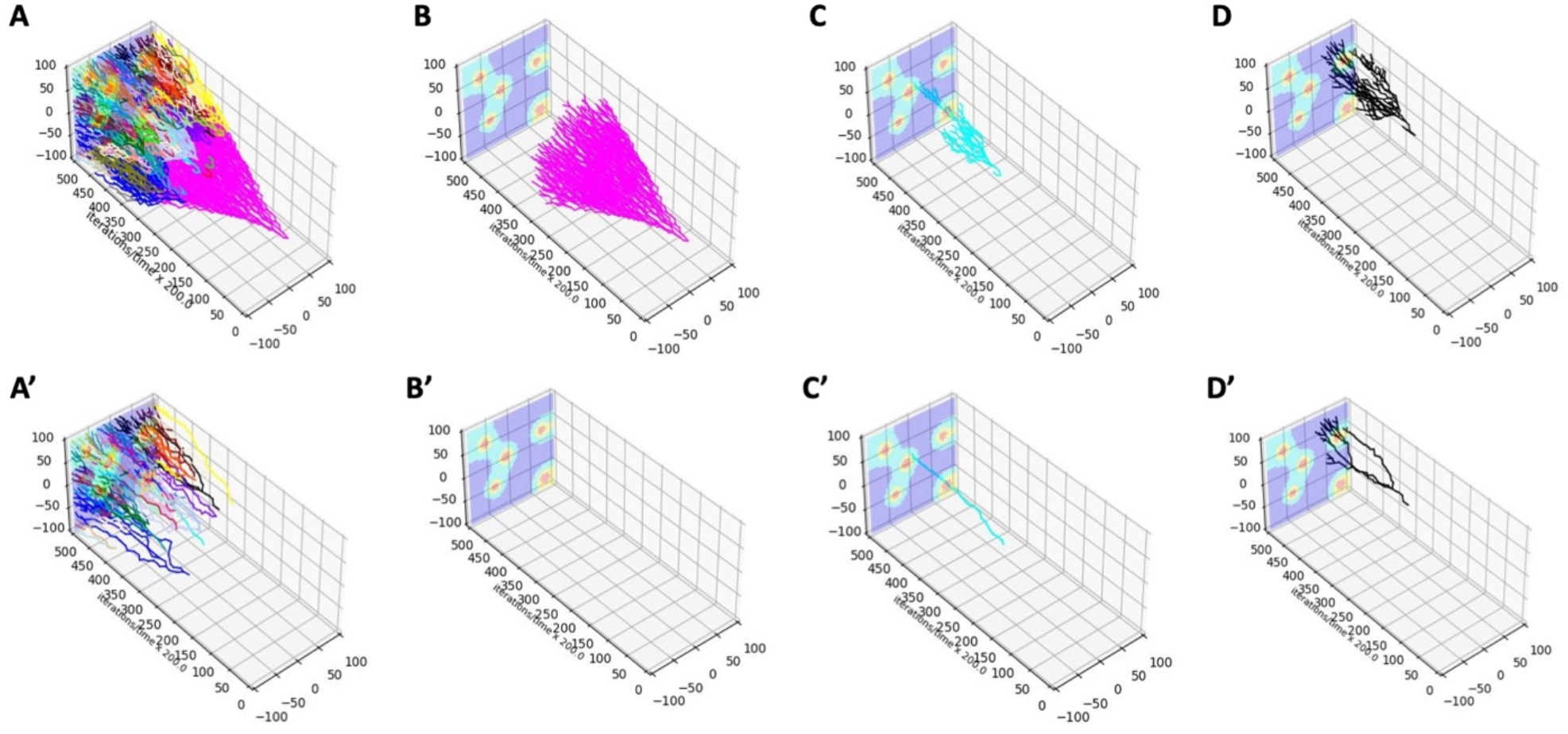
3D lineage trees of individual clones drawn with Python routines. For each clone (denoted by a different color), the top row shows the 3D lineage tree with all cells belonging to that clone (**A** *LinG3DAll* and **B-D** *LinG3DClone* routine), while the bottom row includes only the cells that survived to the end of the simulation (**A’** *LinG3DAliveAll* and **B’-D’** *LinG3DAliveClone* routine). **B-B’:** initial clone #0; **C-C’:** mutated clone #2; **D-D’:** mutated clone #5.

The two presented 3D lineage trees for drug-resistant clones (Fig.7C,D) show how the cells move through the tissue to avoid the areas next to the vessels that contain very high drug concentrations. The surviving cells are localized in the low-drug (blue) regions (Fig.7C’,D’). This information can not be obtained from classical jellyfish evolution graphs (Fig.6G) or phylogenic trees (Fig.6H,J).

To generate these 3D lineage trees, we specified the following function arguments: data directory (`pathData=‘exampleB05’`), spatial domain (`xmin=-100 xmax=100 ymin=-100 ymax=100`), temporal domain (`tmin=0 tmax=10000`), frequency of sampled data (`fileStep=250`), the background indicator (`IsGradient=1`), and the figure saving indicator (`toPrint=1`). For drawing all clones, the total number of clones (`numClones=147`) was specified. For drawing individual clones, the clone numbers (`cloneNum=0,2,5`) were used, respectively.

### Case study of an experimental colony growth assay with identified cellular clones

While our routines were developed to visualize data from computational simulations, they can also be applied to experimental data, provided that data is recorded in the required format. As an illustration, we visualized clones generated during the 2D growth assay. A sequence of frames spaced 12 minutes apart was annotated using in-house MATLAB code to record the coordinates of cells arising from 10 precursor cells chosen from an initial frame. The mother cell-daughter cell relation was also recorded whenever one of the traced cells have divided. The initial frame with ten selected cells is shown in Fig.8A, and the final frame with cells forming the ten clones is shown in Fig.8B. Each clone is drawn with a different color. The full 3D lineage tree of all ten cellular clones (drawn with *LinG3DAll*, MATLAB-based version), is shown in Fig.8C, and four specific 3D lineage trees for individual cellular clones (drawn with *LinG3DClone*, MATLAB-based version) are shown in Fig.8D-G.

**Fig. 8.**
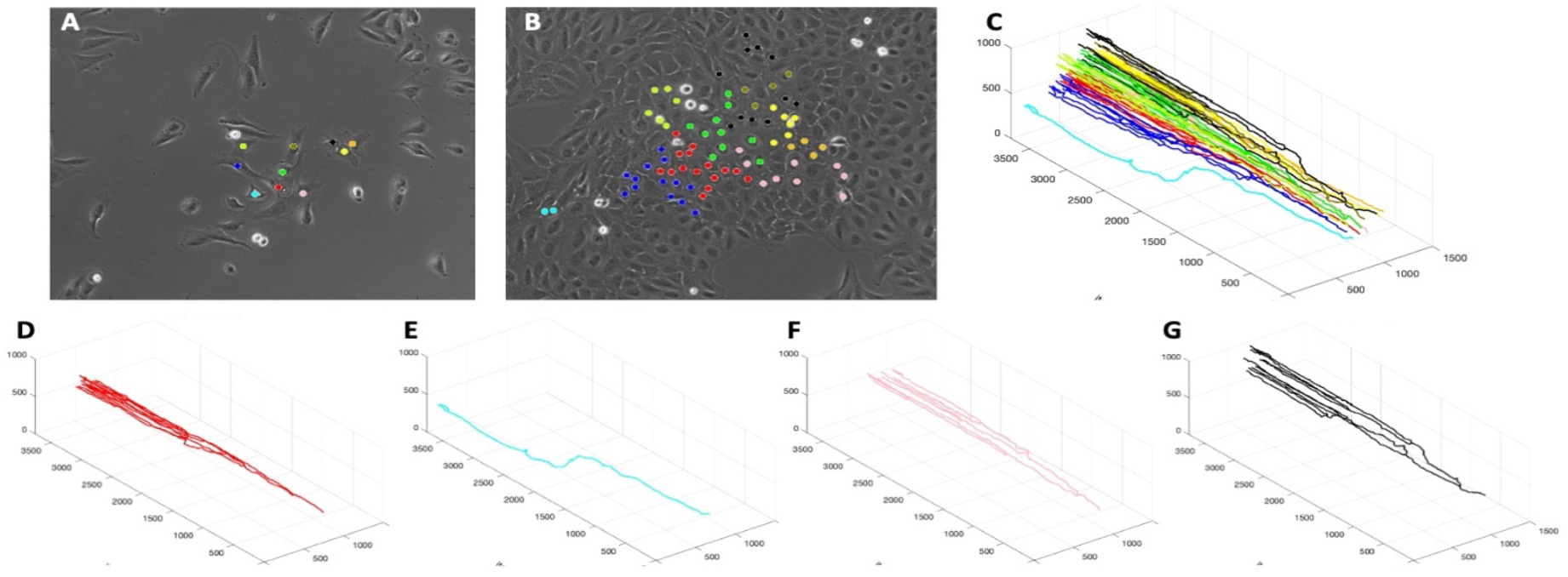
3D lineage trees (MATLAB routines) for individual clones identified in an experimental assay. **A**,**B:** the initial and final frames from a bright field time lapse microscopy of a growing cell colony with 10 different clones indicated by different colors. **C**. 3D lineage trees for all ten cellular clones (*LinG3DAll*); **D-G:** 3D lineage trees for all cells in four selected cellular clones (*LinG3DClone*); each tree color corresponds to cells’ colors in **A** and **B**.

To generate these 3D lineage trees, we had to specify the following input function arguments: **data** directory (`pathData=‘exampleExp’`), spatial domain (`xmin=0 xmax=1500 ymin=0 ymax=1000`), temporal domain (`tmin=0 tmax=864`), frequency of sampled data (`fileStep=4`), the no background indicator (`IsGradient=0`), and the figure saving indicator (`toPrint=1`). For drawing all clones, the total number of clones (`numClones=10`) was specified. For drawing individual clones, the clone numbers (`cloneNum=1,2,10,5`) was used, respectively.

The presented spatio-temporal lineage trees for individual cells revealed that the cell in Fig.8E was quite motile and did not proliferate often, in contrast to other cells (Fig.8D,F,G) that were dividing several times but did not move out at significant distances. These spatial effects are not observed in the classical phylogenic trees or jellyfish evolution graphs.

## Conclusions

The proposed by us *LinG3D* routines generate three-dimensional spatio-temporal trees that show the emergence of cellular clones in time, like classical lineage trees, as well as in space within a patch of the tumor tissue. These routines were designed primarily for application to data generated from simulations of tumor evolution models. However, they can also be applied to experimental cellular clones, if appropriate information can be collected. The routines can illustrate clonal development in growing tumors with or without exposure to anti-cancer treatments. In particular, they can enable analysis of spatial intra-tumor heterogeneity, such as the spatial distribution of cells with specific phenotypes or mutations.

We have previously used the 3D lineage trees routines to inspect tumor areas where drug-induced resistance can arise. These methods allowed us to identify three distinct microenvironmental niches that are prone to the emergence of clones with acquired drug resistance: sanctuaries with low drug exposure, protectorates with medium drug exposure but dense cellularity, and hypoxia niches with low cell proliferation rates (12, 13). Such biological insights would not be recognized without the use of 3D cell lineage trees.

We have shown here that our routines are versatile with respect to the type of data (synthetic or experimental), number of visualized clones (from tens to hundreds), number of visualized timepoints, domain sizes, and computational languages (three popular: R, Python, and MATLAB are included here). The most elaborated 3D lineage tree from our examples is shown in Fig.7A; it took 7 minutes to draw this tree on an Apple laptop computer. While these routines can handle a larger number of clones, the typical 2D jellyfish evolution graphs are applied to cases with the number of clones in tens rather than hundreds, as in the cases we showed in this paper. Future extensions of these routines can include options for changing clone colors in a personalized way, choosing the resolution for background image of microenvironmental factor, or saving the generated images in different formats. Additionally, an option to draw several selected individual clones in the same image could be added.

In summary, visualization of cellular clones in both time and space can provide additional information on whether the particular cells or clones are motile vs. those that are more stationary; whether growing cells spread widely or hover around specific tissue niches. This can help in formulating new research hypotheses.

## Availability of data and materials

All software code and data are available from the following depository: https://github.com/rejniaklab/LinG3D, https://github.com/rejniaklab/pyLinG3D, https://github.com/rejniaklab/r_LinG3D

## Appendix

### Mathematical equations from a model for S1 and S2

To illustrate how the 3D lineage tree algorithms can be applied, we developed a relatively simple agent-based model of the growing tumor exposed to a cytotoxic drug (MATLAB routine: *tumorGrowth_example1.m* and *tumorGrowth_example2.m*). This model is based on our Multi-Cell Lattice-Free, *MultiCell-LF*, framework (12, 14, 15) in which individual cells interact with one another through physical forces and with the surrounding microenvironment via chemical factors, such as drugs or oxygen.

### Computational domain

The model is defined on a square tissue patch: [−100,100] × [−100,100] *μm*^*2*^, that contains five irregularly distributed stationary vessels 𝒱_*i*_ (*i* = 1, … ,5) that are the source of a cytotoxic drug *γ*(x, y, *t*). The tumor cells *X*_*i*_ (*i* = 1, …, 𝒩_*c*_) can proliferate, absorb the drug, undergo random and drug-induced death, and can mutate which results in cell resistance to the drug. The initial condition consists of a single tumor cell located in the middle of the domain with no drug. The no-flux boundary conditions are imposed on the drug domain. The cells are also removed from the system if they move outside of the domain boundaries.

### Drug kinetics

Drug distribution within the tumor tissue depends on three factors: the amount of drug supplied by the vessels, the amount of drug taken up by the tumor cells, and the spatial localization of all blood vessels and cells. The drug *γ*(x, y, *t*) is supplied from each vessel 𝒱_*i*_ with the influx rate 𝒥_*γ*_, it diffuses through the tissue with a constant diffusion coefficient 𝒟_*γ*_, and is absorbed by each cell *X*_*i*_ with the uptake rate 𝒰_*γ*_ proportional to the drug concentration γ(x, y, *t*). The equation of drug kinetics is defined as follows:

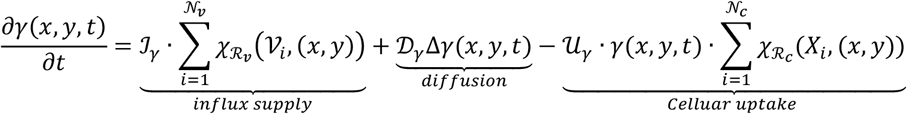

where interactions between the drug grid (x, y) and the individual cells or vessels (*Z* = 𝒱_*i*_ or *X*_*i*_) are specified by the indicator function with radius ℛ:

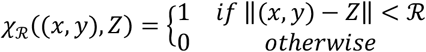

### Cell division and mutation

Each cell is defined by its position *X*_*i*_ and a constant radius ℛ_*c*_. The cell can inspect its vicinity of radius ℛ_*N*,_ and can count the number 𝒩_*i*_ of neighboring cells. Cell age progresses with time 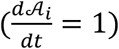, and when the cell reaches maturity, 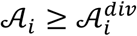, and is not overcrowded, that is the number of cell neighbors does not exceed the prescribed threshold, 𝒩_*i*_ ≤ 𝒩^*max*^, the cell will divide. Upon division of cell *X*_*i*_, two daughter cells 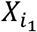 and 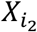 are created. 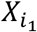 will take position of the mother cell, i.e. 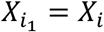 and 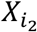 will be located in the vicinity of the mother cell in the randomly selected direction, i.e., 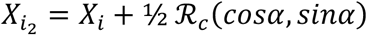, where *α* ∈ [*0*,*2*π] Each daughter cell inherits the mother division age with small noise *ω*, i.e., 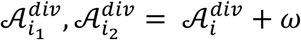, where *ω∈*[−5,5]. The current age for each daughter cell is then set at 0, i.e., 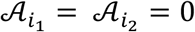.

The mutation status of each daughter cell is inherited from its mother cell, i.e., 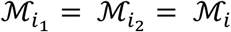. However, each daughter cell can acquire a new mutation status when it is born; the probability of new mutation is determined by 𝒫^*mut*^ (the only parameter we varied in the presented examples). Cell mutation will result in a shorter cell division age, i.e., 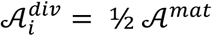, where 𝒜^*mat*^ is the default cell division (maturation) age. It is assumed that the random mutations will start after the cell colony reaches a noticeable size.

### Cell drug absorption, accumulation, and drug-induced death

Each cell absorbs drug *γ*(x, y, *t*) from its immediate vicinity of radius ℛ _*γ*_. The cell uptake rate is 𝒰_*γ*_. Drug accumulation 𝒬_*i*_ by the cell *X*_*i*_ is described by the following equation:

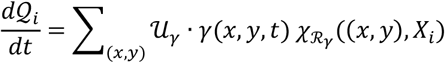

Upon cell division, the levels of absorbed drug for both daughter cells are set to 0, i.e., 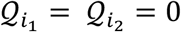. When the drug accumulated by a non-mutated cell (ℳ_*i*_ = *0*) exceeds the prescribed threshold, i.e. 𝒬_*i*_ > 𝒬^*thr*^, the cell will die with a probability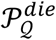. Once the cell dies, it is removed from the system.

All model parameters are listed in Table 1.

**Table 1:** Model physical and computational parameters.

| Parameter | Symbol | Value |
| --- | --- | --- |
| **Physical parameters** |  |  |
| Cell radius | $\mathcal{R}_c$ | $5 \mu m$ |
| Average cell division age | $\mathcal{A}^{mat}$ | 18 min |
| Number of neighbors for overcrowding | $\mathcal{N}^{max}$ | 5 cells |
| Cell neighborhood radius | $\mathcal{R}_N$ | $15 \mu m$ |
| Cell-cell repulsive spring stiffness | $k_c$ | $50 \mu g / \mu m \cdot s^2$ |
| Vessel radius | $\mathcal{R}_v$ | $10 \mu m$ |
| Cell-vessel repulsive spring stiffness | $k_v$ | $100 \mu g / \mu m \cdot s^2$ |
| Medium viscosity | $\nu$ | $250 \mu g / \mu m \cdot s$ |
| Time of drug injection | $\mathcal{N}_{inj}$ | 250 min |
| Drug influx from the vessel | $\mathcal{I}_\gamma$ | $5 \sigma g / \mu m^3 \cdot s$ |
| Drug diffusion coefficient | $\mathcal{D}_\gamma$ | $10 \mu m^2 / s$ |
| Drug uptake rate | $\mathcal{U}_\gamma$ | $0.015 s^{-1}$ |
| Cell sensing radius | $\mathcal{R}_\gamma$ | $3 \mu m$ |
| Drug level for cell killing | $\mathcal{Q}^{thr}$ | $4 \sigma g / \mu m^3$ |
| **Probability parameters** |  |  |
| Probability of mutation | $\mathcal{P}^{mut}$ | 0.05 or 0.005 |
| Probability of random death | $\mathcal{P}^{die}$ | 0.8 |
| Probability of drug induced death | $\mathcal{P}_Q^{die}$ | 0.85 |
| **Computational Parameters** |  |  |
| Domain size | $\Omega$ | $[-100, 100]^2 \mu m$ |
| Grid width | $h_g$ | $5 \mu m$ |
| Time step | $\Delta t$ | $0.25 s$ |
| Number of iterations | $N^{total}$ | 100,000 |
| Scaling parameter | $\sigma$ | $10^{-19}$ |

### Cell random death

All cells, mutated and non-mutated, can undergo random death with a probability 𝒫^*die*^, provided that they are of considerable age, that is their current age 𝒜_*i*_ exceed twice the cell division age 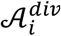. Once the cell dies, it is removed from the system.

### Cell-cell and cell-vessel interactions

Individual cells *X*_*i*_ and *X*_*j*_ can exert a cell-cell repulsive force *f*_*i*,*j*_ (with stiffness *k*_*c*_ and a resting length *2ℛ*_*c*_) to avoid overlapping:

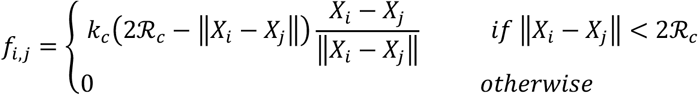

If the cell *X*_*i*_ overlaps with multiple (M) cells, the cumulative repulsive force is:

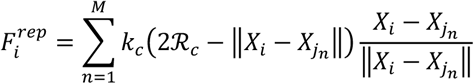

Similarly, to avoid overlapping between cells and vessels, a repulsive force *g*_*i*,*j*_ between the overlapping cell *X*_*i*_ and the vessel, *V*_*j*_ is exerted on the cell:

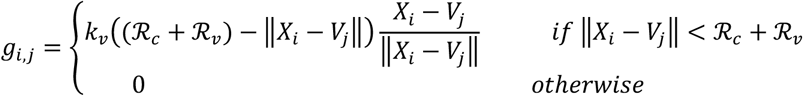

If cell *X*_*i*_ overlaps with multiple (M) vessels, the cumulative repulsive force is

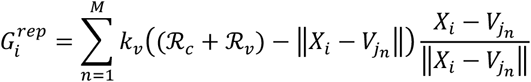

### Cell Relocation

Cell dynamics is governed by Newton’s second law of motion, where the applied forces arise as a result of cell-cell repulsive interactions 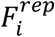, repulsive interactions between cells and vessels 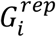, and forces needed to overcome medium viscosity 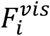, where *m*_*i*_ is cell mass:

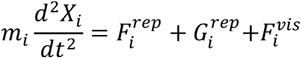

Assuming that springs are overdamped: 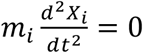, and that viscous force is proportional to cell velocity: 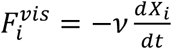, the equation of cell relocation is given by:

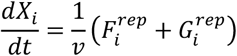

## References

1. Nowell PC. The clonal evolution of tumor cell populations. Science. 1976;194(4260):23–8.

2. Noble R, Burri D, Le Sueur C, Lemant J, Viossat Y, Kather JN, et al. Spatial structure governs the mode of tumour evolution. Nat Ecol Evol. 2021.

3. Miller CA, McMichael J, Dang HX, Maher CA, Ding L, Ley TJ, et al. Visualizing tumor evolution with the fishplot package for R. BMC Genomics. 2016;17(1):880.

4. Gatenbee CD, Schenck RO, Bravo RR, Anderson ARA. EvoFreq: visualization of the Evolutionary Frequencies of sequence and model data. BMC Bioinformatics. 2019;20(1):710.

5. MATLAB. phytree routine documentation and examples. 2021 [Available from: https://www.mathworks.com/help/bioinfo/ref/phytree.html.

6. Yu G. Using ggtree to Visualize Data on Tree-Like Structures. Current Protocols in Bioinformatics 2020;69:e96.

7. Chevenet F, Brun C, Banuls AL, Jacq B, Christen R. TreeDyn: towards dynamic graphics and annotations for analyses of trees. BMC Bioinformatics. 2006;7:439.

8. Murdoch D, Adler D. 3D Visualization Using OpenGL, R package 2022 [Available from: https://CRAN.R-project.org/package=rgl.

9. Blagotic A, Daróczi G. Rapport: a report templating system. R package 2022 [Available from: https://cran.r-project.org/package=rapport.

10. Wickham H, Hester J, Bryan J. readr: Read Rectangular Text Data. R package 2022 [Available from: https://CRAN.R-project.org/package=readr.

11. Wickham H, Hester J, Chang W, J. B. devtools: Tools to Make Developing R Packages Easier. 2022 [Available from: https://devtools.r-lib.org/, https://github.com/r-lib/devtools.

12. Perez-Velazquez J, Rejniak KA. Drug-Induced Resistance in Micrometastases: Analysis of Spatio-Temporal Cell Lineages. Front Physiol. 2020;11:319.

13. Perez-Velazquez J, Gevertz JL, Karolak A, Rejniak KA. Microenvironmental Niches and Sanctuaries: A Route to Acquired Resistance. Adv Exp Med Biol. 2016;936:149–64.

14. Berrouet C, Dorilas N, Rejniak KA, Tuncer N. Comparison of Drug Inhibitory Effects IC50 in Monolayer and Spheroid Cultures. Bull Math Biol. 2020;82(6):68.

15. Kingsley JL, Costello JR, Raghunand N, Rejniak KA. Bridging cell-scale simulations and radiologic images to explain short-time intratumoral oxygen fluctuations. PLoS Comput Biol. 2021;17(7):e1009206.

